# *Paulinella micropora* KR01 holobiont genome assembly for studying primary plastid evolution

**DOI:** 10.1101/794941

**Authors:** Duckhyun Lhee, JunMo Lee, Chung Hyun Cho, Ji-San Ha, Sang Eun Jeong, Che Ok Jeon, Udi Zelzion, Dana C. Price, Ya-Fan Chan, Arwa Gabr, Debashish Bhattacharya, Hwan Su Yoon

## Abstract

The widespread algal and plant (Archaeplastida) plastid originated >1 billion years ago, therefore relatively little can be learned about plastid integration during the initial stages of primary endosymbiosis by studying these highly derived species. Here we focused on a unique model for endosymbiosis research, the photosynthetic amoeba *Paulinella micropora* KR01 (Rhizaria) that underwent a more recent independent primary endosymbiosis about 124 Mya. A total of 149 Gbp of PacBio and 19 Gbp of Illumina data were used to generate the draft assembly that comprises 7,048 contigs with N50=143,028 bp and a total length of 707 Mbp. Genome GC-content was 44% with 76% repetitive sequences. We predicted 32,358 genes that contain 73% of the complete, conserved genes in the BUSCO database. The mean intron length was 882 bp, which is significantly greater than in other Rhizaria (86∼184 bp). Symbiotic bacteria from the culture were isolated and completed genomes were generated from three species (*Mesorhizobium amorphae* Pch-S, *Methylibium petroeiphilum* Pch-M, *Polaromonas* sp. Pch-P) with one draft genome (*Pimelobacter simplex* Pch-N). Our holobiont data establish *P. micropora* KR01 as a model for studying plastid integration and the role of bacterial symbionts in *Paulinella* biology.

## Introduction

Plastids are believed to have originated *via* primary endosymbiosis >1 billion years ago in the ancestor of the Archaeplastida, whereby a cyanobacterium was captured and retained as the photosynthetic organelle (Bhattacharya, et al. 2004; Cenci, et al. 2017; Price, et al. 2012, 2019). The plastid in red and green algae was subsequently engulfed by other eukaryotes *via* secondary or tertiary endosymbiosis. This process gave rise to a vast diversity of extant photosynthetic lineages such as diatoms, dinoflagellates, chlorarachniophytes, and euglenids (Keeling 2010). Despite its great impact on eukaryote evolution and diversity, the understanding of primary endosymbiosis, particularly its early stages, remains limited due to the long span of time that has passed since this event took place (Yoon, et al. 2009). For this reason, the independent and more recent primary endosymbiosis in the rhizarian amoeba, *Paulinella* provides a rare and unique opportunity for studying plastid evolution. In contrast to the highly reduced plastid genomes of algae and plants (i.e., 100-200 kbp in size in Archaeplastida), the cyanobacterium-derived plastid (referred to as the chromatophore) in phototrophic *Paulinella* is in a less derived stage of organelle evolution and is ca. 1 Mbp in size (Lhee, et al. 2017; Nowack, et al. 2008; Reyes-Prieto, et al. 2010; Zhang, et al. 2017). This lineage provides the opportunity to gain fundamental insights into primary endosymbiosis that are not possible by studying Archaeplastida (Qiu, et al. 2012).

As a hallmark of genome erosion, the captured cyanobacterium in *Paulinella* underwent massive gene loss and decrease in GC-content (Lhee, et al. 2019; Nowack, et al. 2008; Qiu, et al. 2012; Reyes-Prieto, et al. 2010). Genes encoded by the previously free-living endosymbiont (most closely related to extant α-cyanobacteria such as *Synechococcus* spp.) were lost outright, retained in the organelle genome, or transferred to host nuclear genome *via* endosymbiotic gene transfer (EGT) (Martin and Herrmann 1998; Reyes-Prieto, et al. 2006). In the case of *Paulinella*, many HGT-derived bacterial genes encode proteins that fill gaps in critical chromatophore pathways and processes (Nowack, et al. 2016). Furthermore, proteins synthesized in the cytoplasm of the amoeba host are transported into the chromatophore where they function in vital processes such as photosynthesis or amino acid biosynthesis (Singer, et al. 2017).

To advance knowledge about primary EGT and genome evolution in photosynthetic *Paulinella*, we generated a draft genome assembly and transcriptome data from *Paulinella micropora* strain KR01 (hereafter, KR01; Lhee, et al. 2017), as well as complete or near-complete draft genomes from four co-cultured bacteria. These data comprise a valuable reference platform for studying the origin and integration of the chromatophore in this independent case of plastid primary origin.

## Materials and Methods

### Sampling, Isolation, and DNA/ RNA Extraction

*Paulinella micropora* KR01 was isolated from a natural freshwater sample collected on August 18, 2009 from Mangae Jeosuji (Reservoir), Chungnam Province, South Korea (see Lhee, et al. 2017). We established an uni-algal, axenic culture of KR01 (for details see Supplementary Information Materials and Methods). The DNeasy Plant Mini Kit (Qiagen, Santa Clarita, Calif.) was used for the DNA extraction and RNA was extracted using the TRI reagent (Molecular Research Center, Cincinnati, USA).

### Sequencing and genome size estimation

To generate the genome data, large fragments of KR01 DNA from the axenic culture were used for PacBio RS II and Illumina HiSeq 2500 sequencing. A bacteria-reduced culture was used for HiSeq 2500 and Illumina Nova Seq 6000 sequencing. For transcriptome sequencing, control and high light stress conditions (control=10 μmol photons m^-2^ sec^-1^; stress=120 μmol photons m^-2^ sec^-1^; see Zhang, et al. 2017) over a 12/ 12h light/ dark cycle and temperature stress conditions (control=24°C; stress=4°C, 38°C) were sequenced using the Illumina Truseq stranded mRNA library prep kit. All sequencing was done by DNA Link Inc. (Seoul, Korea). Detailed methods and the sequencing output are provided in the Supplementary Information Materials and Methods (Supplementary Tables S1 and S2). The 21-mer depth frequency of axenic Nova Seq data was calculated with Jellyfish (Marçais and Kingsford 2011) and the genome size of KR01 was estimated using findGSE (Sun, et al. 2017).

### *De novo* genome assembly and assessment

We generated a total of 149 Gbp of single-molecule real-time (SMRT) sequences using the P6-C4 sequence chemistry with an average subread length of 7.9 kbp. *De novo* assembly was done using the FALCON-Unzip assembler (Chin, et al. 2016) with filtered subreads. The length cut-off option was specified based on the subread N50 value of 12 kbp. The assembly had a contig N50 = 147 kbp for the phased diploid assembly that was “polished” using Quiver (Chin, et al. 2013). We performed error correction using BWA (DePristo, et al. 2011; Li and Durbin 2009) and GATK (DePristo, et al. 2011) with Illumina HiSeq reads to improve the quality of the assembly. Following this procedure, “contaminant” contigs were removed after identification by searching the NCBI bacteria database using BLASTn (Camacho, et al. 2009). Assembled contigs were checked for contamination using BlobTools (Camacho, et al. 2009). A repeat library of the KR01 genome was constructed using RepeatModeler (Smit, et al. 2013-2015) with masking using RepeatMasker (Tarailo-Graovac and Chen 2009)

### Gene prediction and annotation

Gene prediction was done with BRAKER2 (Hoff, et al. 2015; Stanke, et al. 2006). For additional details, see the Supplementary Information Materials and Methods. Predicted genes were annotated using eggNOG-Mapper v2 (Huerta-Cepas, et al. 2017) with the EggNOG database (Huerta-Cepas, et al. 2018). We assessed genome completeness, vis-à-vis the predicted genes, using BUSCO (Simão, et al. 2015). Orthologous group families (OGFs) of proteins were clustered using OrthoFinder (Emms and Kelly 2015).

### Bacteria isolation and genome assembly

See the Supplementary Information Materials and Methods.

## Results and discussion

### KR01 genome assembly and gene prediction

*De novo* assembly using FALCON-Unzip assembler produced 7,440 contigs with N50=147,044 bp with a total length of 763 Mbp (Supplementary Table S3). After removing bacterial and organelle contigs, 7,048 contigs with N50=143,028 bp and a total length of 707 Mbp remained. The genome GC-content was 44% and the genome size was estimated to be 698.345 Mbp using findGSE (Supplementary Figure S1), but the final assembled genome was 707 Mbp in size. Bacterial contamination in the contigs was again tested with BlobTools using GC-content values and read coverage (Supplementary Figure S2). A separate cluster of bacterium-derived contigs was not found using this approach. The KR01 genome contains 76% repetitive sequences (Supplementary Table S4) that were masked prior to protein-coding gene prediction using BRAKER2. Genes with poor transcriptome evidence were removed, resulting in a conservative estimate of 32,358 genes. A total of 89% of these predicted genes (28,791) contained multiple exons (Figure 1) and 3,567 were single exon predictions (Table 1). There was an average of 10 exons per gene. The mean length of introns in KR01 (882 bp) was significantly longer than in other rhizarian species (86∼184 bp). The BUSCO analysis showed 73% complete and 7% fragmented conserved eukaryotic genes (80% total) in the genome. OGF analysis with other Rhizaria identified a core set of 1,807 OGFs in these species. A large number of OGFs (4,842) were shared only between *P. chromatophora* and KR01 (Figure 2).

**Figure 1.**
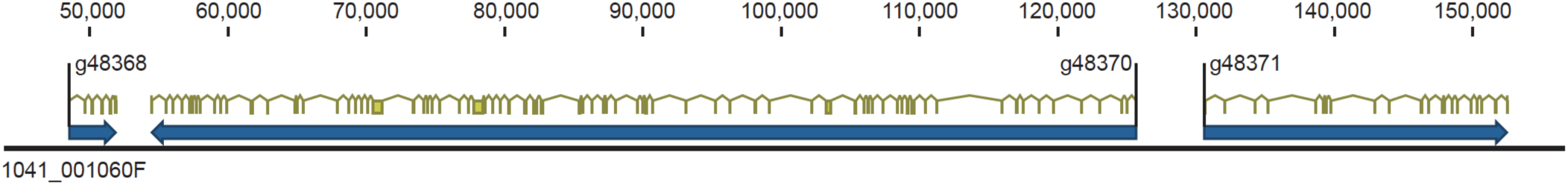
Region of a typical contig (1041_001060F) from the KR01 genome showing the complex intron-exon (filled yellow boxes) structure of many genes (filled blue boxes) in this species. The position in base pairs is shown above the contig.

**Figure 2.**
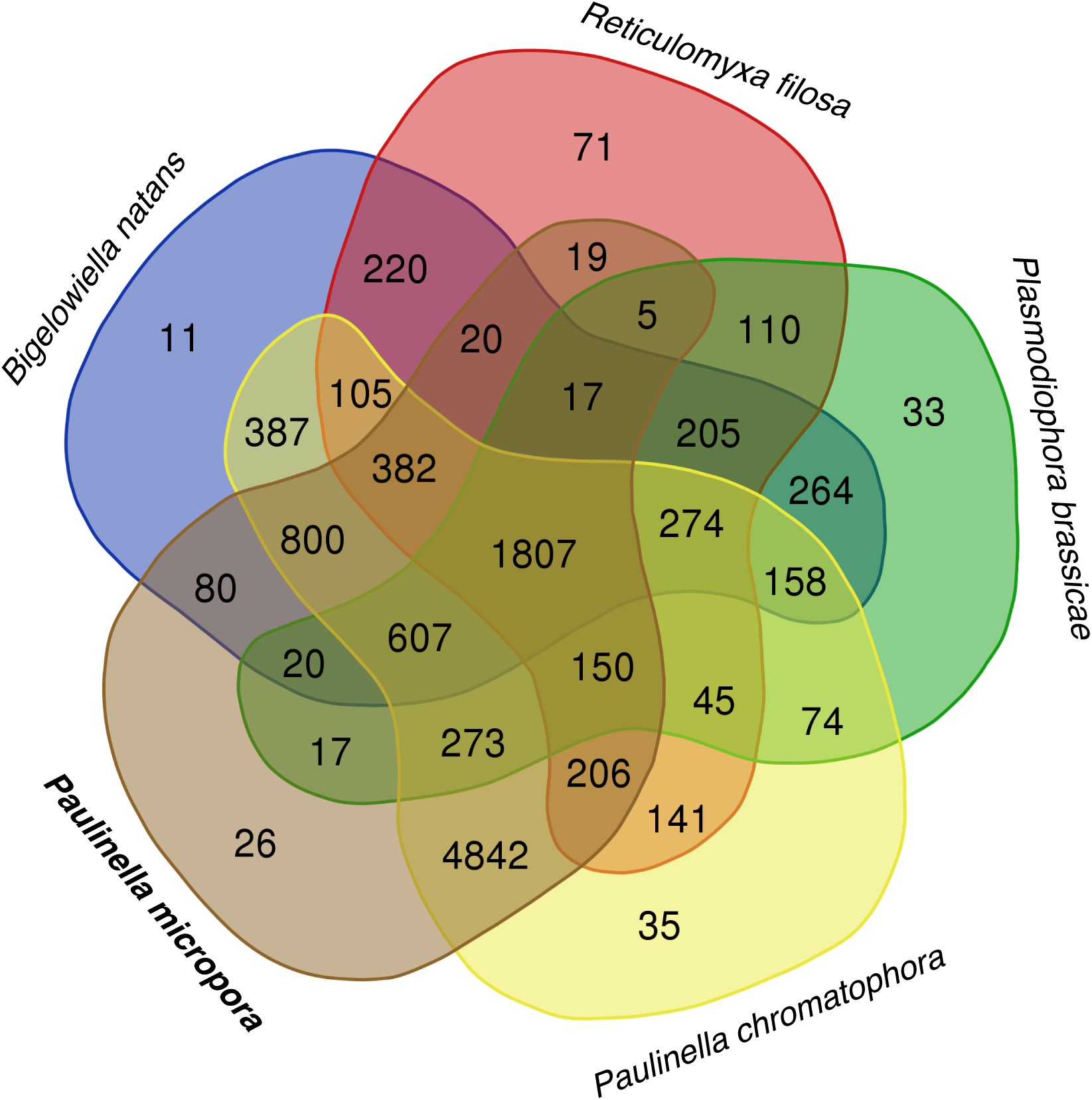
The number of orthologous gene families (OGFs) shared by *Paulinella micropora* KR01 and four other species of Rhizaria species (*Bigelowiella natans, Reticulomyxa filose, Plasmodiophora brassicae*, and *Paulinella chromatophora*).

**Table 1.** Comparison of genome features of available Rhizaria and other photosynthetic species.

|  | *Paulinella micropora* KR01 | *Bigelowiella natans* | *Reticulomyxa filosa* | *Phaeodactylum tricornutum* | *Guillardia theta* | *Chlamydomonas reinhardtii* | *Arabidopsis thaliana* |
| --- | --- | --- | --- | --- | --- | --- | --- |
| Genome size (Mbp) | 707 | 95 | 320 | 27 | 8 | 121 | 140 |
| G-C-content (%) | 44 | 45 | 35 | 49 | 53 | 64 | 36 |
| Protein-coding genes | 32,358 | 21,708 | 40,443 | 10,402 | 24,840 | 15,143 | 26,341 |
| Genes with introns (%) | 89 | 86 | 70 | 47 | 80 | 92 | 79 |
| Mean intron length (bp) | 882 | 184 | 86 | 123 | 110 | 373 | 164 |
| Mean exons per gene | 10 | 9 | 3 | 2 | 6 | 8 | 5 |

Seven different bacteria were isolated from the xenic KR01 culture after streaking on R2A agar (Supplementary Table S5). These bacteria were confirmed by PCR amplification using the universal primer set for 16S rDNA (27F – 1492R). The taxon names we used reflected high similarity in the blast result to known lineages although we used different strain names owing to the isolation from the KR01 culture (Phycosphere of *Paulinella micropora* KR01). The new strain names are as follow: *Pimelobacter simplex* Pch-N, *Bosea thiooxidans* Pch-B, *Mesorhizobium amorphae* Pch-S, *Sphingobium amiense* Pch-SP, *Methylibium petroleiphilum* Pch-M, *Polaromonas* sp. Pch-P and *Pelomonas saccharophila* Pch-PE. The complete genome of three species (*M. amorphae* Pch-S, *M. petroeiphilum* Pch-M, *Polaromonas* sp. Pch-P) and draft genome of one species (*P. simplex* Pch-N) were inferred using the genome data (Supplementary Table S5). We confirmed that the three complete genomes exist in a circular form. The *M. amorphae* Pch-S genome is ca. 7 Mbp in size, has 62% GC-content, and contains 6,613 CDSs, 48 tRNAs, and 6 rRNA genes. The genome size of *M. petroeiphilum* Pch-M is ca. 4 Mbp, with 70% GC-content, and contains 3,814 CDSs, 66 tRNAs, and 3 rRNA genes. The genome of *Polaromonas* sp. is ca. 5 Mbp in size with a 63% GC-contents and contains 4,257 CDSs, 42 tRNAs, and 3 rRNA genes. In contrast to these three complete genomes, the draft genome of *P. simplex* Pch-N is ca. 6 Mbp in size with 73% GC-content, and is comprised of 29 contigs. Gene prediction results show that this strain contains 5,881 CDSs, 47 tRNAs, and 7 rRNA genes. This combination of KR01 genomic and bacterial genome data provide an ideal resource to study the role of bacteria in *Paulinella* holobiont growth and gene function when compared to axenic amoeba cultures.

## Supplementary Materials

Supplementary Tables and figures are available at www.biorxiv.org.

## Data deposition

This project has been deposited at GenBank under the BioProject accession PRJNA568118. The raw sequence data are available at the Marine Genome Information Center (http://www.magic.re.kr/) under the accession numbers MN00067-0001, MN00067-0002 (whole genome) and MN00096-0001, MN00096-0002, MN00126-0001, MN00126-0003, MN00126-0004, MN00126-0005, MN00126-0006, MN00126-0007, MN00126-0008, MN00126-0009, MN00126-0011, MN00126-0013, MN00126-0015, MN00126-0016, MN00126-0003 (for transcriptomes). The bacterial genomes are deposited at GenBank under the accession numbers CP029562 (*M. amorphae* Pch-S), CP029606 (*M. petroeiphilum* Pch-M), CP031013 (*Polaromonas* sp. Pch-P), and WBVM00000000 (*P. simplex* Pch-N).

## Acknowledgements

This study was supported by the Next-generation BioGreen21 Program (PJ01389003) from the RDA (Rural Development Administration) of Korea, the Collaborative Genome Program of the Korea Institute of Marine Science and Technology Promotion (KIMST) funded by the Ministry of Oceans and Fisheries (MOF, 20180430), and the National Research Foundation of Korea (NRF-2017R1A2B3001923) granted to HSY. DB, DCP, and AG were supported by NASA grant 80NSSC19K0462 awarded to DB. YFC was supported by a grant (104-2917-I-564-051) from the Taiwan Ministry of Science and Technology. UZ was supported by the New Jersey Agricultural Experiment Station and the Rutgers Office of Advanced Research Computing. These funding sources were not involved in the conduct of the research and/ or preparation of the article.

## References

Bhattacharya D, Yoon HS, Hackett JD 2004. Photosynthetic eukaryotes unite: endosymbiosis connects the dots. BioEssays 26: 50–60. doi: 10.1002/bies.10376

Camacho C, et al. 2009. BLAST+: architecture and applications. BMC Bioinformatics 10: 421–421. doi: 10.1186/1471-2105-10-421

Cenci U, et al. 2017. Biotic host-pathogen interactions as major drivers of plastid endosymbiosis, Trends Plant Sci 22:316–328. http://dx.doi.org/10.1016/j.tplants.2016.12.007

Chin C-S, et al. 2013. Nonhybrid, finished microbial genome assemblies from long-read SMRT sequencing data. Nature Methods 10: 563. doi: 10.1038/nmeth.2474 https://http://www.nature.com/articles/nmeth.2474 - supplementary-information.

Chin C-S, et al. 2016. Phased diploid genome assembly with single molecule real-time sequencing. bioRxiv. doi: 10.1101/056887

DePristo MA, et al. 2011. A framework for variation discovery and genotyping using next-generation DNA sequencing data. Nature Genetics 43: 491–498. doi: 10.1038/ng.806

Emms DM, Kelly S 2015. OrthoFinder: solving fundamental biases in whole genome comparisons dramatically improves orthogroup inference accuracy. Genome Biol 16: 157. doi: 10.1186/s13059-015-0721-2

Hoff KJ, Lange S, Lomsadze A, Borodovsky M, Stanke M 2015. BRAKER1: unsupervised rna-seq-based genome annotation with GeneMark-ET and AUGUSTUS. Bioinformatics 32: 767–769. doi: 10.1093/bioinformatics/btv661

Huerta-Cepas J, et al. 2017. Fast genome-wide functional annotation through orthology assignment by eggNOG-Mapper. Mol Biol Evol 34: 2115–2122. doi: 10.1093/molbev/msx148

Huerta-Cepas J, et al. 2018. eggNOG 5.0: a hierarchical, functionally and phylogenetically annotated orthology resource based on 5090 organisms and 2502 viruses. Nucleic Acids Research 47: D309–D314. doi: 10.1093/nar/gky1085

Keeling PJ 2010. The endosymbiotic origin, diversification and fate of plastids. Philosophical Transactions of the Royal Society B: Biological Sciences 365: 729–748. doi: doi:10.1098/rstb.2009.0103

Lhee D, et al. 2019. Evolutionary dynamics of the chromatophore genome in three photosynthetic *Paulinella* species. Scientific Reports 9: 2560. doi: 10.1038/s41598-019-38621-8

Lhee D, et al. 2017. Diversity of the photosynthetic *Paulinella* species, with the description of *Paulinella micropora* sp. nov. and the chromatophore genome sequence for strain KR01. Protist 168: 155–170. doi: 10.1016/j.protis.2017.01.003

Li H, Durbin R 2009. Fast and accurate short read alignment with Burrows-Wheeler transform. Bioinformatics (Oxford, England) 25: 1754–1760. doi: 10.1093/bioinformatics/btp324

Marçais G, Kingsford C 2011. A fast, lock-free approach for efficient parallel counting of occurrences of k-mers. Bioinformatics 27: 764–770. doi: 10.1093/bioinformatics/btr011

Martin W, Herrmann RG 1998. Gene transfer from organelles to the nucleus: how much, what happens, and why? Plant Physiology 118: 9. doi: 10.1104/pp.118.1.9

Nowack EC, Melkonian M, Glockner G 2008. Chromatophore genome sequence of *Paulinella* sheds light on acquisition of photosynthesis by eukaryotes. Curr Biol 18: 410–418. doi: 10.1016/j.cub.2008.02.051

Nowack ECM, et al. 2016. Gene transfers from diverse bacteria compensate for reductive genome evolution in the chromatophore of *Paulinella chromatophora*. Proceedings of the National Academy of Sciences of the United States of America 113: 12214–12219. doi: 10.1073/pnas.1608016113

Price DC, et al. 2012. Cyanophora paradoxa genome elucidates origin of photosynthesis in algae and plants. Science 335: 843. doi: 10.1126/science.1213561

Price DC, et al. 2019. Analysis of an improved *Cyanophora paradoxa* genome assembly. DNA Research 26: 287–299. doi: 10.1093/dnares/dsz009

Qiu H, Yang EC, Bhattacharya D, Yoon HS 2012. Ancient gene paralogy may mislead inference of plastid phylogeny. Mol Biol Evol 29: 3333–3343. doi: 10.1093/molbev/mss137

Reyes-Prieto A, Hackett JD, Soares MB, Bonaldo MF, Bhattacharya D 2006. Cyanobacterial contribution to algal nuclear genomes is primarily limited to plastid functions. Current Biology 16: 2320–2325. doi: https://doi.org/10.1016/j.cub.2006.09.063

Reyes-Prieto A, et al. 2010. Differential gene retention in plastids of common recent origin. Mol Biol Evol 27: 1530–1537. doi: 10.1093/molbev/msq032

Simão FA, Waterhouse RM, Ioannidis P, Kriventseva EV, Zdobnov EM 2015. BUSCO: assessing genome assembly and annotation completeness with single-copy orthologs. Bioinformatics 31: 3210–3212. doi: 10.1093/bioinformatics/btv351

Singer A, et al. 2017. Massive protein import into the early-evolutionary-stage photosynthetic organelle of the amoeba *Paulinella chromatophora*. Curr Biol 27: 2763–2773 e2765. doi: 10.1016/j.cub.2017.08.010

Smit A, Hubley R, Green P. 2013-2015. RepeatMasker Open-4.0.

Stanke M, Schöffmann O, Morgenstern B, Waack S 2006. Gene prediction in eukaryotes with a generalized hidden Markov model that uses hints from external sources. BMC Bioinformatics 7: 62–62. doi: 10.1186/1471-2105-7-62

Sun H, Ding J, Piednoël M, Schneeberger K. 2017. findGSE: estimating genome size variation within human and *Arabidopsis* using k-mer frequencies. Bioinformatics 34: 550–557. doi: 10.1093/bioinformatics/btx637

Tarailo-Graovac M, Chen N. 2009. Using RepeatMasker to identify repetitive elements in genomic sequences. Current Protocols in Bioinformatics 25: 4.10.11-14.10.14. doi: 10.1002/0471250953.bi0410s25

Yoon HS, et al. 2009. A single origin of the photosynthetic organelle in different *Paulinella* lineages. BMC Evol Biol 9: 98. doi: 10.1186/1471-2148-9-98

Zhang R, Nowack EC, Price DC, Bhattacharya D, Grossman AR. 2017. Impact of light intensity and quality on chromatophore and nuclear gene expression in *Paulinella* chromatophora, an amoeba with nascent photosynthetic organelles. Plant J 90:221–234. doi: 10.1111/tpj.13488.

