## Supplementary Information for "*Paulinella micropora* KR01 holobiont genome assembly for studying primary plastid evolution"

### Materials and Methods

#### Sampling and isolation

*Paulinella micropora* strain KR01 was isolated from a natural freshwater sample collected on August 18, 2009 from Mangae Jeosuji (Reservoir), Chungnam Province, South Korea (see Lhee, et al. 2017). Cells were found on either the shoreline sediment of the freshwater reservoir or on the surface of aquatic plants that were growing below the water surface. Using capillary pipettes, individual cells were isolated and washed several times with sterilized DY-V medium. After cell isolation, single cells were inoculated to DY-V medium at 20°C with a 12/12 h light/dark cycle with cool-white fluorescent lights. Because photosynthetic *Paulinella* cells generally live in low-light conditions, we used a low light intensity (i.e., 10 µmol photons $m^{-2}$ $s^{-1}$). An axenic culture of *P. micropora* KR01 was produced following the method of (Nowack, et al. 2016). *P. micropora* cells were washed several times using 10 micro meter pore size membrane filter and sprayed on DY-V agar plates. After 3–5 d on the solid medium, bacterial growth was visualized by the formation of whitish halos surrounding nearly all of the *P. micropora* cells. Bacteria-free single cells (halo absent) were picked from the solid medium with a microcapillary and each cell was transferred into DV-V medium. The putatively axenic state of cultures was checked microscopically and total genomic DNA was sequenced with Illumina HiSeq 2500. Assembled contigs with 16S rDNA were matched to the *P. micropora* chromatophore 16S rDNA nucleotide sequence using Blastn. No other bacterial 16S rDNA sequences were found.

#### PacBio library construction and sequencing

Using the Covaris G-tube, we generated 20 kbp fragments by shearing genomic DNA according to the manufacturer's protocol. We used the AMpureXP bead purification system to remove small fragments. A total of 5 µg for each sample was used as input into library preparation. The SMRTbell library was constructed using the SMRTbell™ Template Prep Kit 1.0 (PN 100-259-100). Using the BluePippin Size selection system, we removed the small fragments to allow generation of large-insert libraries. After a sequencing primer was annealed to the SMRTbell template, DNA polymerase was bound to the complex (DNA/Polymerase Binding kit P6). Following the polymerase binding reaction, the MagBead Kit was used to bind the library complex with MagBeads before sequencing. MagBead bound complexes provide for more reads per SMRT Cell. This polymerase-SMRTbell-adaptor complex was then loaded into zero-mode waveguides (ZMWs). The SMRTbell library was sequenced using 115 SMRT cells (Pacific Biosciences) using C4 chemistry (DNA sequencing Reagent 4.0) and 1 × 240 minute movies were captured for each SMRT cell using the PacBio RS (Pacific Biosciences) sequencing platform.

**Illumina Hi-Seq library construction and sequencing**

DNA/ RNA purity was determined by analyzing 1 µl of total DNA/ RNA extract on a NanoDrop8000 spectrophotometer. Total RNA integrity was checked using an Agilent Technologies 2100 Bioanalyzer with an RNA Integrity Number (RIN) value.

DNA sequencing libraries were prepared according to the manufacturer's instructions (Truseq DNA PCR-Free Library Prep kit). The mRNA sequencing libraries were prepared according to the manufacturer's instructions (Illumina Truseq stranded mRNA library prep kit) and the mRNA was purified and fragmented from total RNA (1 µg) using poly-T oligo-attached magnetic beads with two rounds of purification. Cleaved RNA fragments primed with random hexamers were reverse transcribed into first strand cDNA using reverse transcriptase, random primers, and dUTP in place of dTTP. The incorporation of dUTP quenches the second strand during amplification, because the polymerase does not incorporate past this nucleotide. These cDNA fragments then had the addition of a single 'A' base and subsequent ligation of the adapter. The products were purified and enriched with PCR to create the final strand specific cDNA libraries. The quality of the amplified libraries was verified by capillary electrophoresis (Bioanalyzer, Agilent).

After QPCR using SYBR Green PCR Master Mix (Applied Biosystems), we combined libraries that index tagged in equimolar amounts in the pool. Cluster generation occurred in the flow cell on the cBot automated cluster generation system (Illumina). The flow cell was loaded on the HiSeq 2500 sequencing system (Illumina) with 2 x 100bp reagents.

**Illumina NovaSeq 6000 library construction and sequencing**

Input DNA for sequencing needs to be as intact as possible, with an OD 260/280 ratio of 1.8–2.0. We checked the quality of DNA by 1% agarose gel electrophoresis and the Qubit dsDNA HS Assay Kit (Thermo Fisher Scientific). The DNA library was prepared according to Illumina Truseq Nano DNA Library prep protocol. For sample library preparation, 0.2 µg for insert 550 bp size of high molecular weight genomic DNA was randomly sheared to yield DNA fragments using the Covaris S2 system. The fragments were made blunt-ended and phosphorylated, and a single 'A' nucleotide was added to the 3'-termini of the fragments in preparation for ligation to an adapter that has a single-base 'T' overhang. Adapter ligation at both ends of the genomic DNA fragment conferred different sequences at the 5' and 3' ends of each strand in the genomic fragment. Ligated DNA was PCR amplified to enrich for fragments that have adapters on both ends. The quality of the amplified libraries was verified by capillary electrophoresis (Bioanalyzer, Agilent). After QPCR using the SYBR Green PCR Master Mix (Applied Biosystems), we combined libraries that were index tagged in equimolar amounts in the pool. WGS sequencing is performed using an Illumina NovaSeq 6000 system following provided protocols for 2 x 100 bp reagents.

**Gene prediction and annotation**

Gene prediction was done with BRAKER2 (Hoff, et al. 2015; Stanke, et al. 2006). The following data were used for hints: RNA-seq data (high and control light condition 0h, 0.5h, 6h, 12h, 18h, 30h, 42h), do novo assembled RNA-Seq data using Trinity (Grabherr, et al. 2011), and *Paulinella chromatophora* CCAC0185 protein data (Nowack, et al. 2016). To filter out genes with low RNA-Seq read coverage, axenic RNA-Seq data (different temperature: 24°C, 4°C, and 38°C) were aligned to the genome assembly using the STAR tool with default parameters (Dobin, et al. 2013). Predicted genes with average coverage less than one were removed. Among the different transcripts from same gene, we choose the first transcript (named ".t1" in the gtf output file) and removed the remainder.

**Bacteria isolation and genome construction**

To isolate bacteria, culture medium supernatant was spread on R2A agar plates and incubated at room temperature. After 3~5 days, bacterial colonies were isolated and transferred to new R2A agar for purification. Using this method, seven different species were isolated and identified by 16S rDNA PCR and sequencing (27F – 1492R). Genomic DNA from the cells was extracted using the Wizard Genomic DNA Purification kit (Promega, USA), following the protocol recommended by the manufacturer, and sequenced using a combination of PacBio RSII single-molecule real-time (SMRT) sequencing at DNA Link (Seoul, Korea). *De novo* assembly of the sequencing reads, derived from the PacBio SMRT sequencing, was done using the hierarchical genome assembly process (HGAP). Bacterial genome was annotated using Prokka (Seemann 2014), and the quality check was done using Checkm (Parks, et al. 2015).

### Reference

Dobin A, et al. 2013. STAR: ultrafast universal RNA-seq aligner. Bioinformatics (Oxford, England) 29: 15-21. doi: 10.1093/bioinformatics/bts635

Grabherr MG, et al. 2011. Full-length transcriptome assembly from RNA-Seq data without a reference genome. Nature Biotechnology 29: 644. doi: 10.1038/nbt.1883

https://http://www.nature.com/articles/nbt.1883 - supplementary-information

Hoff KJ, Lange S, Lomsadze A, Borodovsky M, Stanke M 2015. BRAKER1: Unsupervised RNA-Seq-based genome annotation with GeneMark-ET and AUGUSTUS. Bioinformatics 32: 767-769. doi: 10.1093/bioinformatics/btv661

Nowack ECM, et al. 2016. Gene transfers from diverse bacteria compensate for reductive genome evolution in the chromatophore of *Paulinella chromatophora*. Proceedings of the National Academy of Sciences of the United States of America 113: 12214-12219. doi: 10.1073/pnas.1608016113

Parks DH, Imelfort M, Skennerton CT, Hugenholtz P, Tyson GW 2015. CheckM: assessing the quality of microbial genomes recovered from isolates, single cells, and metagenomes. Genome Research 25: 1043-1055. doi: 10.1101/gr.186072.114

Seemann T 2014. Prokka: rapid prokaryotic genome annotation. Bioinformatics 30: 2068-2069. doi: 10.1093/bioinformatics/btu153

Stanke M, Schöffmann O, Morgenstern B, Waack S 2006. Gene prediction in eukaryotes with a generalized hidden Markov model that uses hints from external sources. BMC Bioinformatics 7: 62-62. doi: 10.1186/1471-2105-7-62

**Supplementary Figure S1.** Frequency distribution of the 21-mer graph used to estimate the genome size of *Paulinella micropora* strain KR01.

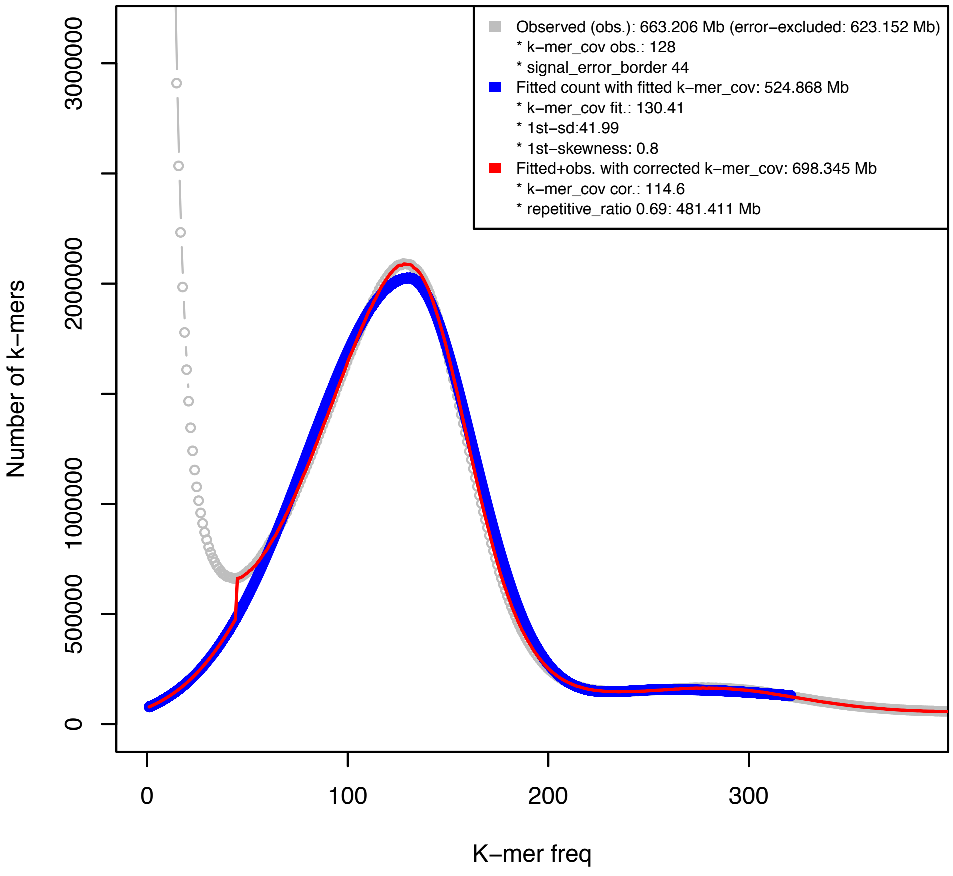

**Supplementary Figure S2.** BlobPlot of the assembled KR01 contigs. Sequences in the assembly are depicted as circles, with the diameter scaled proportional to sequence length and colored by taxonomic annotation (at the rank of 'order') based on BLASTn and Diamond blastx similarity search results provided in this order and using taxrule 'bestsumorder'. The circles are positioned on the X-axis based on their GC-content and on the Y-axis based on the sum of coverage.

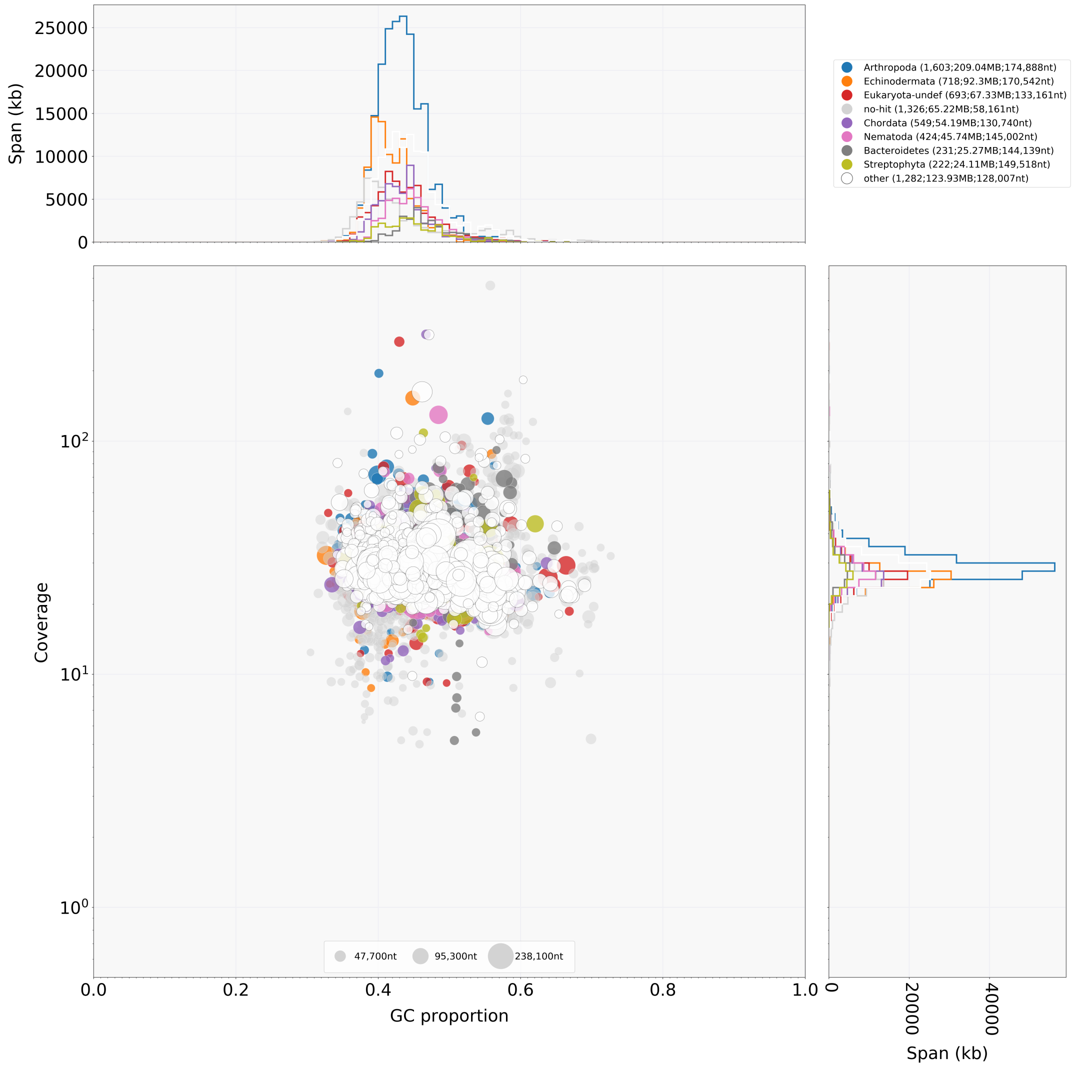

**Supplementary Table S1.** Genomic sequencing information for *Paulinella micropora* KR01.

| PacBio RS sequencing subreads summary | | | | |
|---|---|---|---|---|
| **Sample** | **Number of Bases** | **Number of Reads** | **N50 Read Length** | **Mean Read Length** |
| *Paulinella micropora* strain KR01 xenic culture | 148,962,628,479 | 18,796,456 | 11,938 | 7,925 |

| Illumina HiSeq 2500 sequencing statistics | | | | | | |
|---|---|---|---|---|---|---|
| **Sample ID** | **Read Order** | **Index** | **Yield(Bases)** | **Reads** | **% of >= Q30 Bases(PF)** | **Mean Quality Score(PF)** |
| *Paulinella micropora* strain KR01 xenic culture | 1 | ATCACG | 6,469,415,721 | 64,053,621 | 86.8 | 34.7 |
| *Paulinella micropora* strain KR01 xenic culture | 2 | ATCACG | 6,469,415,721 | 64,053,621 | 81.65 | 33.46 |
| *Paulinella micropora* strain KR01 axenic culture | 1 | GTTTCG | 12,511,206,431 | 123,873,331 | 80.01 | 33.24 |
| *Paulinella micropora* strain KR01 axenic culture | 2 | GTTTCG | 12,511,206,431 | 123,873,331 | 71.7 | 31.29 |

| Illumina NovaSeq 6000 sequencing statistics | | | | | | |
|---|---|---|---|---|---|---|
| **Sample ID** | **Read Order** | **Index** | **Yield(Bases)** | **Reads** | **% of >= Q30 Bases(PF)** | **Mean Quality Score(PF)** |
| *Paulinella micropora* strain KR01 axenic culture | R1 | GTTTCG | 96,625,765,144 | 956,690,744 | 90.48 | 34.33 |
| *Paulinella micropora* strain KR01 axenic culture | R2 | GTTTCG | 96,625,765,144 | 956,690,744 | 81.28 | 32.52 |

**Supplementary Table S2.** RNA sequencing information for *Paulinella micropora* KR01.

| Illumina HiSeq 2500 sequencing statistics | | | | | | | |
|---|---|---|---|---|---|---|---|
| **Sample ID** | **Condition** | **Read Order** | **Index** | **Yield(Bases)** | **Reads** | **% of >= Q30 Bases(PF)** | **Mean Quality Score(PF)** |
| *Paulinella micropora* strain KR01 xenic culture | Control light condition time point 0h in 12/12h(light/dark) cycle | 1 | CGATGT | 2,631,106,863 | 26,050,563 | 90 | 35.52 |
| *Paulinella micropora* strain KR01 xenic culture | Control light condition time point 0h in 12/12h(light/dark) cycle | 2 | CGATGT | 2,631,106,863 | 26,050,563 | 84.89 | 34.32 |
| *Paulinella micropora* strain KR01 xenic culture | Control light condition time point 6h in 12/12h(light/dark) cycle | 1 | TGACCA | 2,238,166,565 | 22,160,065 | 89.79 | 35.47 |
| *Paulinella micropora* strain KR01 xenic culture | Control light condition time point 6h in 12/12h(light/dark) cycle | 2 | TGACCA | 2,238,166,565 | 22,160,065 | 85.47 | 34.44 |
| *Paulinella micropora* strain KR01 xenic culture | Control light condition time point 12h in 12/12h(light/dark) cycle | 1 | ACAGTG | 1,674,610,300 | 16,580,300 | 90 | 35.52 |
| *Paulinella micropora* strain KR01 xenic culture | Control light condition time point 12h in 12/12h(light/dark) cycle | 2 | ACAGTG | 1,674,610,300 | 16,580,300 | 85.13 | 34.39 |
| *Paulinella micropora* strain KR01 xenic culture | Control light condition time point 18h in 12/12h(light/dark) cycle | 1 | CTTGTA | 1,589,326,506 | 15,735,906 | 89.98 | 35.52 |
| *Paulinella micropora* strain KR01 xenic culture | Control light condition time point 18h in 12/12h(light/dark) cycle | 2 | CTTGTA | 1,589,326,506 | 15,735,906 | 85.28 | 34.42 |
| *Paulinella micropora* strain KR01 xenic culture | Control light condition time point 24.5h in 12/12h(light/dark) cycle | 1 | GCCAAT | 1,723,048,284 | 17,059,884 | 89.78 | 35.47 |
| *Paulinella micropora* strain KR01 xenic culture | Control light condition time point 24.5h in 12/12h(light/dark) cycle | 2 | GCCAAT | 1,723,048,284 | 17,059,884 | 85.16 | 34.39 |
| *Paulinella micropora* strain KR01 xenic culture | Control light condition time point 26h in 12/12h(light/dark) cycle | 1 | CAGATC | 1,627,572,681 | 16,114,581 | 89.82 | 35.47 |
| *Paulinella micropora* strain KR01 xenic culture | Control light condition time point 26h in 12/12h(light/dark) cycle | 2 | CAGATC | 1,627,572,681 | 16,114,581 | 84.99 | 34.34 |
| *Paulinella micropora* strain KR01 xenic culture | Control light condition time point 0h in 12/12h(light/dark) cycle, replicate 1 | 1 | GTTTCG | 2528536414 | 25035014 | 96.01 | 36.78 |
| *Paulinella micropora* strain KR01 xenic culture | Control light condition time point 0h in 12/12h(light/dark) cycle, replicate 1 | 2 | GTTTCG | 2528536414 | 25035014 | 93.89 | 36.3 |
| *Paulinella micropora* strain KR01 xenic culture | Control light condition time point 0h in 12/12h(light/dark) cycle, replicate 2 | 1 | GCCAAT | 3316105124 | 32832724 | 93.24 | 36.14 |
| *Paulinella micropora* strain KR01 xenic culture | Control light condition time point 0h in 12/12h(light/dark) cycle, replicate 2 | 2 | GCCAAT | 3316105124 | 32832724 | 90.63 | 35.56 |
| *Paulinella micropora* strain KR01 xenic culture | Control light condition time point 0h in 12/12h(light/dark) cycle, replicate 3 | 1 | GGCTAC | 2432450569 | 24083669 | 94.83 | 36.5 |

| | | | | | | | |
|---|---|---|---|---|---|---|---|
| *Paulinella micropora* strain KR01 xenic culture | Control light condition time point 0h in 12/12h(light/dark) cycle, replicate 3 | 2 | GGCTAC | 2432450569 | 24083669 | 92.42 | 35.96 |
| *Paulinella micropora* strain KR01 xenic culture | Control light condition time point 0.5h in 12/12h(light/dark) cycle, replicate 1 | 1 | ACTTGA | 2580633123 | 25550823 | 94.4 | 36.41 |
| *Paulinella micropora* strain KR01 xenic culture | Control light condition time point 0.5h in 12/12h(light/dark) cycle, replicate 1 | 2 | ACTTGA | 2580633123 | 25550823 | 92.09 | 35.9 |
| *Paulinella micropora* strain KR01 xenic culture | Control light condition time point 0.5h in 12/12h(light/dark) cycle, replicate 2 | 1 | CAGATC | 2740588439 | 27134539 | 95.89 | 36.74 |
| *Paulinella micropora* strain KR01 xenic culture | Control light condition time point 0.5h in 12/12h(light/dark) cycle, replicate 2 | 2 | CAGATC | 2740588439 | 27134539 | 93.5 | 36.18 |
| *Paulinella micropora* strain KR01 xenic culture | Control light condition time point 0.5h in 12/12h(light/dark) cycle, replicate 3 | 1 | TGACCA | 2975130033 | 29456733 | 93.27 | 36.15 |
| *Paulinella micropora* strain KR01 xenic culture | Control light condition time point 0.5h in 12/12h(light/dark) cycle, replicate 3 | 2 | TGACCA | 2975130033 | 29456733 | 91 | 35.63 |
| *Paulinella micropora* strain KR01 xenic culture | Control light condition time point 6h in 12/12h(light/dark) cycle, replicate 1 | 1 | GTGGCC | 2818888790 | 27909790 | 95.95 | 36.76 |
| *Paulinella micropora* strain KR01 xenic culture | Control light condition time point 6h in 12/12h(light/dark) cycle, replicate 1 | 2 | GTGGCC | 2818888790 | 27909790 | 94.11 | 36.35 |
| *Paulinella micropora* strain KR01 xenic culture | Control light condition time point 6h in 12/12h(light/dark) cycle, replicate 2 | 1 | AGTTCC | 2952432505 | 29232005 | 93.4 | 36.18 |
| *Paulinella micropora* strain KR01 xenic culture | Control light condition time point 6h in 12/12h(light/dark) cycle, replicate 2 | 2 | AGTTCC | 2952432505 | 29232005 | 90.72 | 35.58 |
| *Paulinella micropora* strain KR01 xenic culture | Control light condition time point 6h in 12/12h(light/dark) cycle, replicate 3 | 1 | GTTTCG | 2694965022 | 26682822 | 94.97 | 36.54 |
| *Paulinella micropora* strain KR01 xenic culture | Control light condition time point 6h in 12/12h(light/dark) cycle, replicate 3 | 2 | GTTTCG | 2694965022 | 26682822 | 92.51 | 35.98 |
| *Paulinella micropora* strain KR01 xenic culture | Control light condition time point 12h in 12/12h(light/dark) cycle, replicate 1 | 1 | CTTGTA | 2751819336 | 27245736 | 93.5 | 36.2 |
| *Paulinella micropora* strain KR01 xenic culture | Control light condition time point 12h in 12/12h(light/dark) cycle, replicate 1 | 2 | CTTGTA | 2751819336 | 27245736 | 90.17 | 35.45 |
| *Paulinella micropora* strain KR01 xenic culture | Control light condition time point 12h in 12/12h(light/dark) cycle, replicate 2 | 1 | CGTACG | 2899486285 | 28707785 | 95 | 36.55 |
| *Paulinella micropora* strain KR01 xenic culture | Control light condition time point 12h in 12/12h(light/dark) cycle, replicate 2 | 2 | CGTACG | 2899486285 | 28707785 | 92.6 | 36.01 |
| *Paulinella micropora* strain KR01 xenic culture | Control light condition time point 12h in 12/12h(light/dark) cycle, replicate 3 | 1 | GAGTGG | 3114988066 | 30841466 | 95.3 | 36.62 |
| *Paulinella micropora* strain KR01 xenic culture | Control light condition time point 12h in 12/12h(light/dark) cycle, replicate 3 | 2 | GAGTGG | 3114988066 | 30841466 | 92.16 | 35.92 |

| | | | | | | | |
|---|---|---|---|---|---|---|---|
| *Paulinella micropora* strain KR01 xenic culture | Control light condition time point 18h in 12/12h(light/dark) cycle, replicate 1 | 1 | CTTGTA | 2835141205 | 28070705 | 95.72 | 35.72 |
| *Paulinella micropora* strain KR01 xenic culture | Control light condition time point 18h in 12/12h(light/dark) cycle, replicate 1 | 2 | CTTGTA | 2835141205 | 28070705 | 93.2 | 35.15 |
| *Paulinella micropora* strain KR01 xenic culture | Control light condition time point 18h in 12/12h(light/dark) cycle, replicate 2 | 1 | CGATGT | 3663647235 | 36273735 | 93.46 | 36.2 |
| *Paulinella micropora* strain KR01 xenic culture | Control light condition time point 18h in 12/12h(light/dark) cycle, replicate 2 | 2 | CGATGT | 3663647235 | 36273735 | 89.99 | 35.4 |
| *Paulinella micropora* strain KR01 xenic culture | Control light condition time point 18h in 12/12h(light/dark) cycle, replicate 3 | 1 | ACTGAT | 3272703404 | 32403004 | 93.28 | 36.15 |
| *Paulinella micropora* strain KR01 xenic culture | Control light condition time point 18h in 12/12h(light/dark) cycle, replicate 3 | 2 | ACTGAT | 3272703404 | 32403004 | 90.37 | 35.49 |
| *Paulinella micropora* strain KR01 xenic culture | Control light condition time point 30h in 12/12h(light/dark) cycle, replicate 1 | 1 | GGCTAC | 2603080878 | 25773078 | 94.16 | 36.35 |
| *Paulinella micropora* strain KR01 xenic culture | Control light condition time point 30h in 12/12h(light/dark) cycle, replicate 1 | 2 | GGCTAC | 2603080878 | 25773078 | 91.58 | 35.77 |
| *Paulinella micropora* strain KR01 xenic culture | Control light condition time point 30h in 12/12h(light/dark) cycle, replicate 2 | 1 | ATGTCA | 3706578598 | 36698798 | 94.49 | 36.43 |
| *Paulinella micropora* strain KR01 xenic culture | Control light condition time point 30h in 12/12h(light/dark) cycle, replicate 2 | 2 | ATGTCA | 3706578598 | 36698798 | 92.05 | 35.88 |
| *Paulinella micropora* strain KR01 xenic culture | Control light condition time point 30h in 12/12h(light/dark) cycle, replicate 3 | 1 | ATTCCT | 3480729367 | 34462667 | 93.2 | 36.13 |
| *Paulinella micropora* strain KR01 xenic culture | Control light condition time point 30h in 12/12h(light/dark) cycle, replicate 3 | 2 | ATTCCT | 3480729367 | 34462667 | 90.81 | 35.59 |
| *Paulinella micropora* strain KR01 xenic culture | Control light condition time point 42h in 12/12h(light/dark) cycle, replicate 1 | 1 | ACTGAT | 2517612759 | 24926859 | 96.02 | 36.78 |
| *Paulinella micropora* strain KR01 xenic culture | Control light condition time point 42h in 12/12h(light/dark) cycle, replicate 1 | 2 | ACTGAT | 2517612759 | 24926859 | 94.46 | 36.43 |
| *Paulinella micropora* strain KR01 xenic culture | Control light condition time point 42h in 12/12h(light/dark) cycle, replicate 2 | 1 | ATTCCT | 2491865536 | 24671936 | 95.53 | 35.69 |
| *Paulinella micropora* strain KR01 xenic culture | Control light condition time point 42h in 12/12h(light/dark) cycle, replicate 2 | 2 | ATTCCT | 2491865536 | 24671936 | 94.04 | 35.35 |
| *Paulinella micropora* strain KR01 xenic culture | Control light condition time point 42h in 12/12h(light/dark) cycle, replicate 3 | 1 | GATCAG | 3281006513 | 32485213 | 93.99 | 36.31 |
| *Paulinella micropora* strain KR01 xenic culture | Control light condition time point 42h in 12/12h(light/dark) cycle, replicate 3 | 2 | GATCAG | 3281006513 | 32485213 | 90.79 | 35.57 |
| *Paulinella micropora* strain KR01 xenic culture | High light stress light condition time point 0h in 12/12h(light/dark) cycle, replicate 1 | 1 | GAGTGG | 2532558840 | 25074840 | 96.24 | 36.84 |

| | | | | | | | |
|---|---|---|---|---|---|---|---|
| *Paulinella micropora* strain KR01 xenic culture | High light stress light condition time point 0h in 12/12h(light/dark) cycle, replicate 1 | 2 | GAGTGG | 2532558840 | 25074840 | 93.6 | 36.24 |
| *Paulinella micropora* strain KR01 xenic culture | High light stress light condition time point 0h in 12/12h(light/dark) cycle, replicate 2 | 1 | TTAGGC | 2442119602 | 24179402 | 94.28 | 36.38 |
| *Paulinella micropora* strain KR01 xenic culture | High light stress light condition time point 0h in 12/12h(light/dark) cycle, replicate 2 | 2 | TTAGGC | 2442119602 | 24179402 | 92.19 | 35.92 |
| *Paulinella micropora* strain KR01 xenic culture | High light stress light condition time point 0h in 12/12h(light/dark) cycle, replicate 3 | 1 | TTAGGC | 3392632723 | 33590423 | 94.87 | 36.51 |
| *Paulinella micropora* strain KR01 xenic culture | High light stress light condition time point 0h in 12/12h(light/dark) cycle, replicate 3 | 2 | TTAGGC | 3392632723 | 33590423 | 92.97 | 36.08 |
| *Paulinella micropora* strain KR01 xenic culture | High light stress light condition time point 0.5h in 12/12h(light/dark) cycle, replicate 1 | 1 | TAGCTT | 3049219694 | 30190294 | 94.4 | 36.41 |
| *Paulinella micropora* strain KR01 xenic culture | High light stress light condition time point 0.5h in 12/12h(light/dark) cycle, replicate 1 | 2 | TAGCTT | 3049219694 | 30190294 | 92.19 | 35.91 |
| *Paulinella micropora* strain KR01 xenic culture | High light stress light condition time point 0.5h in 12/12h(light/dark) cycle, replicate 2 | 1 | ATCACG | 2507103406 | 24822806 | 94.08 | 36.33 |
| *Paulinella micropora* strain KR01 xenic culture | High light stress light condition time point 0.5h in 12/12h(light/dark) cycle, replicate 2 | 2 | ATCACG | 2507103406 | 24822806 | 91.91 | 35.85 |
| *Paulinella micropora* strain KR01 xenic culture | High light stress light condition time point 0.5h in 12/12h(light/dark) cycle, replicate 3 | 1 | ATCACG | 2788959561 | 27613461 | 94.68 | 36.46 |
| *Paulinella micropora* strain KR01 xenic culture | High light stress light condition time point 0.5h in 12/12h(light/dark) cycle, replicate 3 | 2 | ATCACG | 2788959561 | 27613461 | 92.76 | 36.03 |
| *Paulinella micropora* strain KR01 xenic culture | High light stress light condition time point 6h in 12/12h(light/dark) cycle, replicate 1 | 1 | AGTCAA | 2814060586 | 27861986 | 93.28 | 36.15 |
| *Paulinella micropora* strain KR01 xenic culture | High light stress light condition time point 6h in 12/12h(light/dark) cycle, replicate 1 | 2 | AGTCAA | 2814060586 | 27861986 | 90.9 | 35.62 |
| *Paulinella micropora* strain KR01 xenic culture | High light stress light condition time point 6h in 12/12h(light/dark) cycle, replicate 2 | 1 | ACTTGA | 3524682648 | 34897848 | 94.96 | 36.53 |
| *Paulinella micropora* strain KR01 xenic culture | High light stress light condition time point 6h in 12/12h(light/dark) cycle, replicate 2 | 2 | ACTTGA | 3524682648 | 34897848 | 92.88 | 36.06 |
| *Paulinella micropora* strain KR01 xenic culture | High light stress light condition time point 6h in 12/12h(light/dark) cycle, replicate 3 | 1 | CGATGT | 2847440884 | 28192484 | 96.09 | 36.8 |
| *Paulinella micropora* strain KR01 xenic culture | High light stress light condition time point 6h in 12/12h(light/dark) cycle, replicate 3 | 2 | CGATGT | 2847440884 | 28192484 | 93.57 | 36.21 |
| *Paulinella micropora* strain KR01 xenic culture | High light stress light condition time point 12h in 12/12h(light/dark) cycle, replicate 1 | 1 | CAGATC | 3575292738 | 35398938 | 93.37 | 36.17 |
| *Paulinella micropora* strain KR01 xenic culture | High light stress light condition time point 12h in 12/12h(light/dark) cycle, replicate 1 | 2 | CAGATC | 3575292738 | 35398938 | 90.34 | 35.47 |

| | | | | | | | |
|---|---|---|---|---|---|---|---|
| *Paulinella micropora* strain KR01 xenic culture | High light stress light condition time point 12h in 12/12h(light/dark) cycle, replicate 2 | 1 | GATCAG | 3107906148 | 30771348 | 94.93 | 36.53 |
| *Paulinella micropora* strain KR01 xenic culture | High light stress light condition time point 12h in 12/12h(light/dark) cycle, replicate 2 | 2 | GATCAG | 3107906148 | 30771348 | 92.34 | 35.94 |
| *Paulinella micropora* strain KR01 xenic culture | High light stress light condition time point 12h in 12/12h(light/dark) cycle, replicate 3 | 1 | TGACCA | 2692175705 | 26655205 | 95.95 | 36.76 |
| *Paulinella micropora* strain KR01 xenic culture | High light stress light condition time point 12h in 12/12h(light/dark) cycle, replicate 3 | 2 | TGACCA | 2692175705 | 26655205 | 94.14 | 36.35 |
| *Paulinella micropora* strain KR01 xenic culture | High light stress light condition time point 18h in 12/12h(light/dark) cycle, replicate 1 | 1 | GCCAAT | 2607821515 | 25820015 | 95.96 | 36.76 |
| *Paulinella micropora* strain KR01 xenic culture | High light stress light condition time point 18h in 12/12h(light/dark) cycle, replicate 1 | 2 | GCCAAT | 2607821515 | 25820015 | 94.13 | 36.35 |
| *Paulinella micropora* strain KR01 xenic culture | High light stress light condition time point 18h in 12/12h(light/dark) cycle, replicate 2 | 1 | TAGCTT | 2458794904 | 24344504 | 95.02 | 36.55 |
| *Paulinella micropora* strain KR01 xenic culture | High light stress light condition time point 18h in 12/12h(light/dark) cycle, replicate 2 | 2 | TAGCTT | 2458794904 | 24344504 | 93.02 | 36.09 |
| *Paulinella micropora* strain KR01 xenic culture | High light stress light condition time point 18h in 12/12h(light/dark) cycle, replicate 3 | 1 | ACAGTG | 2605201676 | 25794076 | 96.05 | 36.79 |
| *Paulinella micropora* strain KR01 xenic culture | High light stress light condition time point 18h in 12/12h(light/dark) cycle, replicate 3 | 2 | ACAGTG | 2605201676 | 25794076 | 93.93 | 36.31 |
| *Paulinella micropora* strain KR01 xenic culture | High light stress light condition time point 30h in 12/12h(light/dark) cycle, replicate 1 | 1 | CGTACG | 2622765576 | 25967976 | 95.98 | 36.77 |
| *Paulinella micropora* strain KR01 xenic culture | High light stress light condition time point 30h in 12/12h(light/dark) cycle, replicate 1 | 2 | CGTACG | 2622765576 | 25967976 | 93.87 | 36.29 |
| *Paulinella micropora* strain KR01 xenic culture | High light stress light condition time point 30h in 12/12h(light/dark) cycle, replicate 2 | 1 | GTGAAA | 3387106710 | 33535710 | 94.1 | 36.34 |
| *Paulinella micropora* strain KR01 xenic culture | High light stress light condition time point 30h in 12/12h(light/dark) cycle, replicate 2 | 2 | GTGAAA | 3387106710 | 33535710 | 91.7 | 35.8 |
| *Paulinella micropora* strain KR01 xenic culture | High light stress light condition time point 30h in 12/12h(light/dark) cycle, replicate 3 | 1 | GTGGCC | 2514886163 | 24899863 | 94.83 | 36.5 |
| *Paulinella micropora* strain KR01 xenic culture | High light stress light condition time point 30h in 12/12h(light/dark) cycle, replicate 3 | 2 | GTGGCC | 2514886163 | 24899863 | 92.73 | 36.03 |
| *Paulinella micropora* strain KR01 xenic culture | High light stress light condition time point 42h in 12/12h(light/dark) cycle, replicate 1 | 1 | CCGTCC | 3288862495 | 32562995 | 93.93 | 36.3 |
| *Paulinella micropora* strain KR01 xenic culture | High light stress light condition time point 42h in 12/12h(light/dark) cycle, replicate 1 | 2 | CCGTCC | 3288862495 | 32562995 | 91.66 | 35.78 |
| *Paulinella micropora* strain KR01 xenic culture | High light stress light condition time point 42h in 12/12h(light/dark) cycle, replicate 2 | 1 | GTCCGC | 3080914201 | 30504101 | 94.02 | 36.32 |

| | | | | | | | |
|---|---|---|---|---|---|---|---|
| *Paulinella micropora* strain KR01 xenic culture | High light stress light condition time point 42h in 12/12h(light/dark) cycle, replicate 2 | 2 | GTCCGC | 3080914201 | 30504101 | 91.98 | 35.87 |
| *Paulinella micropora* strain KR01 xenic culture | High light stress light condition time point 42h in 12/12h(light/dark) cycle, replicate 3 | 1 | ACAGTG | 2925562162 | 28965962 | 93.4 | 36.18 |
| *Paulinella micropora* strain KR01 xenic culture | High light stress light condition time point 42h in 12/12h(light/dark) cycle, replicate 3 | 2 | ACAGTG | 2925562162 | 28965962 | 90.48 | 35.52 |
| *Paulinella micropora* strain KR01 axenic culture | Normal growing condition temperature 24C during light cycle | 1 | TGACCA | 2,853,341,809 | 28,250,909 | 93.61 | 36.22 |
| *Paulinella micropora* strain KR01 axenic culture | Normal growing condition temperature 24C during light cycle | 2 | TGACCA | 2,853,341,809 | 28,250,909 | 91.03 | 35.54 |
| *Paulinella micropora* strain KR01 axenic culture | 180min in 4C during light cycle | 1 | CGATGT | 2,898,432,653 | 28,697,353 | 93.72 | 36.25 |
| *Paulinella micropora* strain KR01 axenic culture | 180min in 4C during light cycle | 2 | CGATGT | 2,898,432,653 | 28,697,353 | 90.35 | 35.36 |
| *Paulinella micropora* strain KR01 axenic culture | 30min in 38C during light cycle | 1 | CTTGTA | 2,562,882,070 | 25,375,070 | 94.16 | 36.33 |
| *Paulinella micropora* strain KR01 axenic culture | 30min in 38C during light cycle | 2 | CTTGTA | 2,562,882,070 | 25,375,070 | 92.32 | 35.83 |

**Supplementary Table S3.** Statistics of the assembled genome of *Paulinella micropora* KR01.

| | unzip | | quiver | illumina error correction | After contamination filtering |
|---|---|---|---|---|---|
| | primary | haplotig | primary | primary | primary |
| Number of contigs | 7,489 | 14,359 | 7,440 | 7,440 | 7,048 |
| Total size of contigs | 758,466,000 | 442,730,074 | 763,065,834 | 762,995,855 | 707124085 |
| Longest contig size | 3,532,458 | 7,447,058 | 3,535,397 | 3,535,397 | 952128 |
| Number of contigs > 1K nt | 7,440 | 14,340 | 7,440 | 7,440 | 7,048 |
| Number of contigs > 10K nt | 7,438 | 12,947 | 7,438 | 7,438 | 7,046 |
| Number of contigs > 100K nt | 2,630 | 122 | 2,652 | 2,651 | 2,539 |
| Number of contigs > 1M nt | 8 | 5 | 8 | 8 | 0 |
| Mean contig size | 101,277 | 30,833 | 102,563 | 102,553 | 100,330 |
| Median contig size | 71,741 | 24,304 | 72,935 | 72,936 | 73,826 |
| N50 contig length | 146,131 | 37,568 | 147,053 | 147,044 | 143028 |
| L50 contig count | 1,555 | 3,491 | 1,556 | 1,556 | |
| GC Contents (%) | 44.9 | 45.19 | 44.92 | 44.91 | 43.77 |
| WGS Mapping Rate (%) | 89.11% | | | | |
| Properly Paired (%) | 82.05% | | | | |

**Supplementary Table S4.** Detailed information about repeat content in the *Paulinella micropora* KR01 genome.

| | number of elements* | length occupied | percentage of sequence |
|---|---|---|---|
| SINEs: | 2469 | 394164 bp | 0.06% |
| ALUs | 0 | 0 bp | 0.00% |
| MIRs | 0 | 0 bp | 0.00% |
| LINEs: | 111130 | 39550603 bp | 5.59% |
| LINE1 | 704 | 576375 bp | 0.08% |
| LINE2 | 84958 | 18953155 bp | 2.68% |
| L3/CR1 | 776 | 1387677 bp | 0.20% |
| LTR elements: | 94513 | 73322144 bp | 10.37% |
| ERVL | 0 | 0 bp | 0.00% |
| ERVL-MaLRs | 0 | 0 bp | 0.00% |
| ERV_classI | 967 | 50694 bp | 0.01% |
| ERV_classII | 102 | 203170 bp | 0.03% |
| DNA elements: | 214482 | 68469204 bp | 9.68% |
| hAT-Charlie | 8512 | 2357537 bp | 0.33% |
| TcMar-Tigger | 0 | 0 bp | 0.00% |
| Unclassified: | 479029 | 205283536 bp | 29.03% |
| Total interspersed repeats: | | 387019651 bp | 54.73% |
| Small RNA: | 2469 | 394164 bp | 0.06% |
| Satellites: | 20466 | 2232404 bp | 0.32% |
| Simple repeats: | 1142803 | 150007754 bp | 21.21% |
| Low complexity: | 39459 | 5065862 bp | 0.72% |

**Supplementary Table S5.** Features associated with the bacteria isolated from a xenic culture of *Paulinella micropora* KR01.

| **No.** | **Isolated bacteria name** | **Genome feature** | | | | | | | | | | | |
|---|---|---|---|---|---|---|---|---|---|---|---|---|---|
| | | **Geome construction** | **contig no. for identification** | **Contig No.** | **N50 (bp)** | **Total length (Mb)** | **Gene No.** | **CDS No.** | **tRNA** | **rRNA** | **GC contents** | **Competeness (%)** | **Contamination (%)** |
| **1** | ***Pimelobacter simplex* Pch-N** | Draft_genome (NCBI_uploaded) | contig_10 | 22 | 3,389,742 | 5.8 | 5,935 | 5,881 | 47 | 7 | 72.8% | 99.22 | 1.73 |
| **2** | ***Bosea thiooxidans* Pch-B** | - | x (PCR confirm) | - | - | - | - | - | | | - | | |
| **3** | ***Mesorhizobium amorphae* Pch-S** | Complete (NCBI_released) | contig.1.cir | 1 (Circular form) | - | 6.6 | 6,332 | 6,276 | 46 | 6 | 62.1% | 99.59 | 1.65 |
| **4** | ***Pelomonas saccharophila* Pch-PE** | - | (PCR confirm) | - | - | - | - | - | | | - | | |
| **5** | ***Sphingobium amiense* Pch-SP** | - | x (PCR confirm) | - | - | - | - | - | | | - | | |
| **6** | ***Methylibium petroleiphilum* Pch-M** | Complete (NCBI_released) | contig.1.cir | 1 (Circular form) | - | 3.9 | 3,761 | 3,689 | 65 | 3 | 69.6% | 99.84 | 0.86 |
| **7** | ***Polaromonas sp.* Pch-P** | Complete (NCBI_released) | contig.1.cir | 1 (Circular form) | - | 4.5 | 4,228 | 4,179 | 42 | 3 | 63% | 99.41 | 0.19 |

| **No.** | **Isolated bacteria name** | **NCBI accession number** | **Description** |
|---|---|---|---|
| **1** | ***Pimelobacter simplex* Pch-N** | WBVM00000000 | - Steroid dehydrogenation, 2,4-DNP degradation, histamine degradation activity (Sukhodolskaya et al., 2017) |
| **2** | ***Bosea thiooxidans* Pch-B** | - | - N-fixing ability (Rivas et al., 2007) (but isolated M. amophae doesn't have this gene set)<br>- Residing in the root nodule (sami et al., 2016) |
| **3** | ***Mesorhizobium amorphae* Pch-S** | CP029562 | - N-fixing ability (Rivas et al., 2007) (but isolated M. amophae doesn't have this gene set)<br>- Residing in the root nodule (sami et al., 2016) |
| **4** | ***Pelomonas saccharophila* Pch-PE** | - | '- One of bacteria consortium units in Cyanobactera phycosphere (Velichko et al., 2015) |
| **5** | ***Sphingobium amiense* Pch-SP** | - | - Nonylphenol-degrading bacterium (ref. Yuuji et al., 2003)<br>- Nitrogenase activity (Marnie et al., 2009)<br>- IAA production (Marnie at al., 2013) |
| **6** | ***Methylibium petroleiphilum* Pch-M** | CP029606 | - Methyl tert-butyl ether (MTBE) is used as a carbon source in this strain (Kane, S. R. et al. 2001) |
| **7** | ***Polaromonas sp.* Pch-P** | CP031013 | - there are few discription |
